# Cell identity revealed by precise cell cycle state mapping links data modalities

**DOI:** 10.1101/2024.09.04.610488

**Authors:** Saeed Alahmari, Andrew Schultz, Jordan Albrecht, Vural Tagal, Zaid Siddiqui, Sandhya Prabhakaran, Issam El Naqa, Alexander Anderson, Laura Heiser, Noemi Andor

**Affiliations:** Department of Computer Science, Najran University, Najran 66462, Saudi Arabia; Department of Integrated Mathematical Oncology, H Lee Moffitt Cancer Center and Research Institute, Tampa, FL, USA; Department of Radiation Oncology, Baylor College of Medicine, Houston, TX, USA; Department of Machine Learning, H Lee Moffitt Cancer Center and Research Institute, Tampa, FL, USA; Department of Biomedical Engineering, Oregon Health and Science University, Portland, OR, USA

## Abstract

Several methods for cell cycle inference from sequencing data exist and are widely adopted. In contrast, methods for classification of cell cycle state from imaging data are scarce. We have for the first time integrated sequencing and imaging derived cell cycle pseudo-times for assigning 449 imaged cells to 693 sequenced cells at an average resolution of 3.4 and 2.4 cells for sequencing and imaging data respectively. Data integration revealed thousands of pathways and organelle features that are correlated with each other, including several previously known interactions and novel associations. The ability to assign the transcriptome state of a profiled cell to its closest living relative, which is still actively growing and expanding opens the door for genotype-phenotype mapping at single cell resolution forward in time.

## 1 Introduction

A cell’s transcriptome is a channel of information propagation; it is a snapshot of how a cell interacts with and responds to its environment. Cells co-existing in the same tumor, or in the same cell line [1, 2], often differ in their genomes and transcriptomes. We and others have shown that these differences often correlate with morphological and structural differences between cells [3, 4] and several imaging- and transcriptome-derived feature pairs have been identified. Expression of cell membrane- and cell surface genes can inform the size of a cell [5, 6, 7, 8]. The copy number of mitochondrial DNA (mtDNA) can serve as proxy for inter-cell differences in the number of mitochondria they carry and several regulators of cell shape and morphology have been identified [9, 10], including: FLO11, STE2 [5], ELN, and TGFB1 [11].

New microfluidic platforms have been designed to link phenotypic analysis of living cells to single-cell sequencing [12, 13]. However, the throughput of these platforms is limited to a few hundred cells, precluding learning from these data, general rules that link fitness to transcriptome snapshots. Yuan et al. developed SCOPE-seq – a microwell array based platform, and claims that a more aggressive cell loading of the platform could increase throughput to several thousands of cells [14]. Nevertheless, all solutions available so far that link phenotypic and genomic measurement, have done so over a *narrow time window* : typically less than one cell generation.

We propose *in-silico* mapping of sequenced and imaged cells as a solution to extend the temporal reach of transcriptome-phenotype integration. We leverage the influence cell cycle progression has on a cell’s transcriptome, morphology and subcellular organization to integrate transcriptome profiles obtained from scRNA-seq of a stomach cancer cell line (NCI-N87) [15] with 3D brightfield images from the same cell line. This article focuses on a prerequisite for integrating sequencing and imaging data: inferring a cell’s position along the cell cycle continuum from 3D images. Our article is structured as follows: we first evaluate whether the Fluorescent Ubiquitination-based Cell Cycle Indicator (FUCCI) can inform cell cycle progression at a higher temporal resolution than simply distinguishing G1 and S/G2/M phases of the cell cycle. We then use a previously developed convolutional neural network (CNN) [16], to calculate the spatial coordinates of nucleus, mitochondria and cytoplasm [17] in each imaged cell. Next we use size- and shape statistics calculated for these subcellular compartments to assign cells to a continuum of cell cycle progression. We conclude with an outlook on how the assigned cell cycle state can be used to link imaged cells to sequenced cells.

Projecting a snapshot of the transcriptome onto subcellular architecture is a natural way to integrate information on the pathway membership of genes and their localization into a holistic view of a cell.

## 2 Results

### Continuous temporal resolution on cell cycle progression with FUCCI

We established NCI-N87 cells stably transfected with FUCCI-vector plasmids (Methods 4.1). We acquired 3D images of the transfected cells on a Leica TCS SP8 equipped with an oil-immersion objective (Methods 4.2) on brightfield and green/red fluorescence channels (Figure 1A). Green cells were classified as cells in G1 phase and comprised 30% of the population. Red cells and double positive (yellow) cells were classified as cells in the S phase (9.5% cells) and G2/M phase, respectively (32% of the population). Cells with no color were assigned as G1/S transitional and comprised 28% of the population (Figure 1B,C).

**Figure 1.**
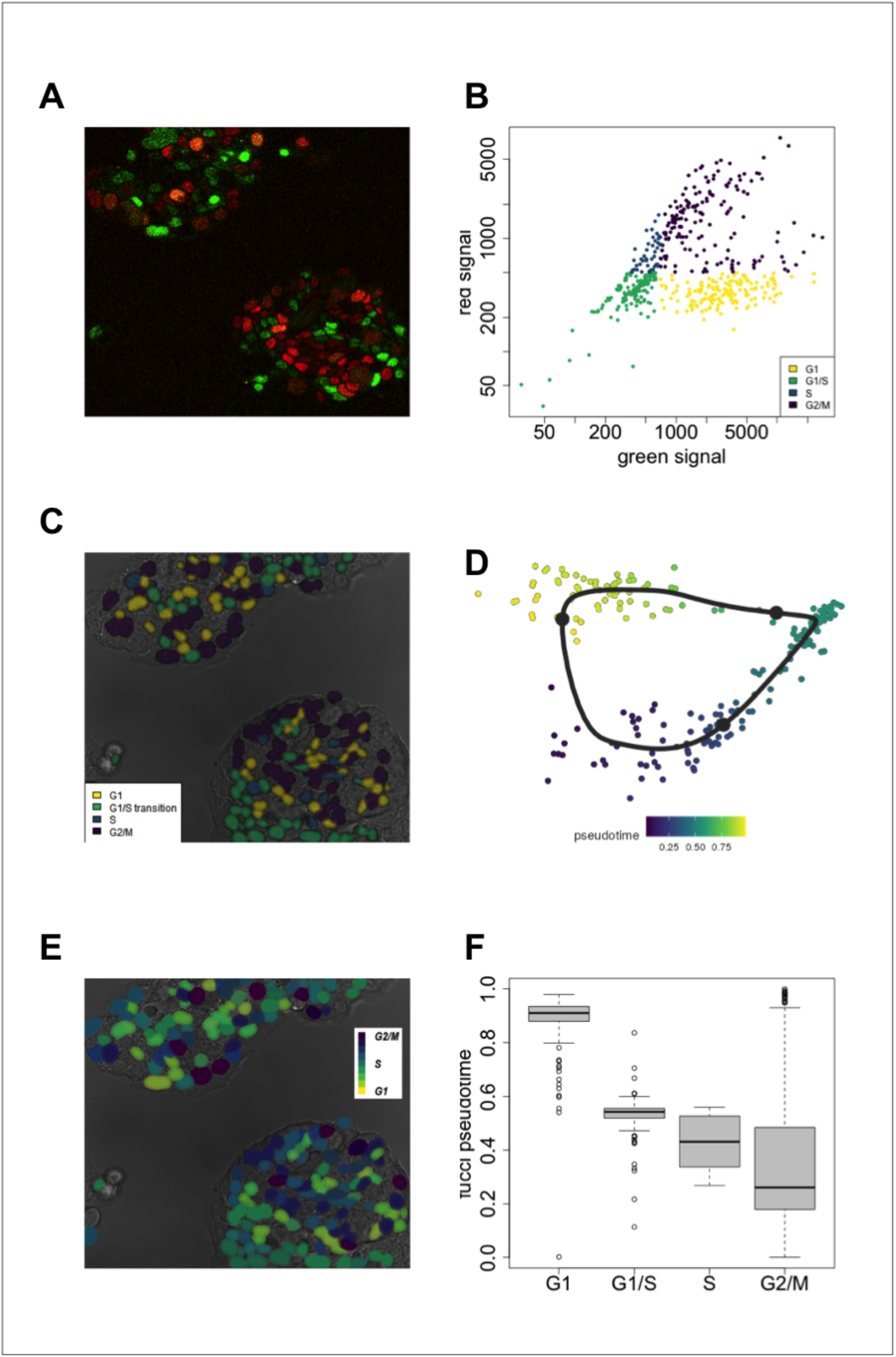
FUCCI to characterize discrete and continuous cell cycle progression in NCI-N87. A. Green and red Fucci channels overlayed. B,C: Fucci based discrete cell cycle classification. D: Fucci based pseudotime inference quantifies continuous cell cycle progression. E: Fucci based pseudotime trajectory in PCA space. Every dot represents a cell. F: Pseudotime is differentially distributed across the four discrete cell cycle classes. Data and code to produce this figure can be found on the project GitHub repository (Methods 4).

Each cell cycle phase has been described as a series of steps that proceeds at a fixed rate [18, 19, 20]. Biologically, the steps refer to a sequence of events that need to be completed for the cell to proceed to the next cell cycle phase (e.g. accumulation of a molecular factor [21, 22] or degradation of proteins [23]). We asked whether FUCCI could quantify cell cycle progression at a temporal resolution higher than that given by distinction of the four cell cycle phases. Specifically, we asked whether FUCCI reporters combined with 3D imaging can rank cells according to how many of the steps within a given cell cycle phase each cell has completed. We calculated 45 intensity features for each cell across the three imaging channels and projected them onto PCA space as input to the Angle method for inferring pseudotime trajectories (Figure 1D). This method computes the angle with respect to the origin in a two-dimensional PCA space and uses this angle as pseudotime for generating a cyclical trajectory [24]. The inferred pseudotime was correlated with and differentially distributed across the four discrete cell cycle classes (Figure 1E,F).

These results indicate that unsupervised methods can be applied to FUCCI based features to widen the application of FUCCI to continuous cell cycle mapping.

### Mitochondrial, cytoplasmic and nuclear changes inform cell cycle progressiom

Despite having revolutionized live-cell imaging of cell cycle transitions, imaging FUCCI labeled cells requires excitation light, which can cause photobleaching and phototoxicity [25]. We therefore asked if a similar continuous cell cycle mapping could be achieved with label-free imaging.

We used 3D images containing the nuclei, mitochondria or cytoplasm of NCI-N87 cells (Methods 4.1), to train a previously developed label-free U-Net convolutional neural network (CNN) [16]. U-Net is an encoder-decoder artificial neural network proposed by Ronneberger et al. for semantic segmentation [26]. Models were trained to predict the coordinates of nucleus, mitochondria and cytoplasm respectively (Methods 4.3). Model performance was evaluated using the Pearson correlation coefficient between the pixel intensities of the model’s predicted output and the independent fluorescence images, with all three signals scoring well above background (Pearson *r >* 0.71 for signal vs. Pearson *r <* 0.04 for background; Wilcoxon rank sum test *P <* 0.002; Fig. 2). The segmentation of nuclei was done using Cellpose [27] – a general purpose algorithm for cell segmentation (Methods 4.3), while mitochondrial and cytoplasmic signals were assigned to nuclei with DBSCAN [28] (Methods 4.10).

**Figure 2.**
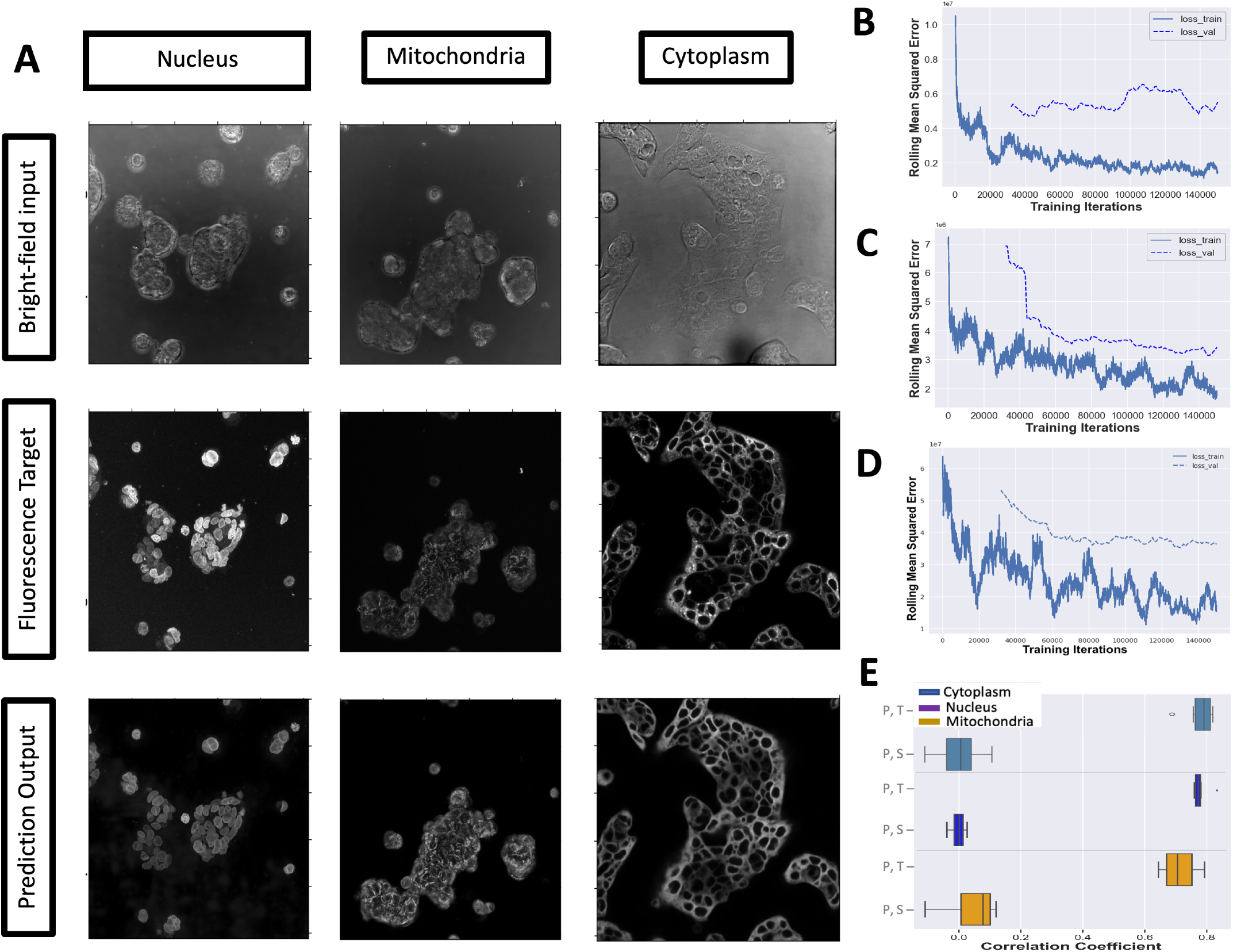
Evaluating performance of a CNN to predict the spatial coordinates of nuclei, mitochondria and cytoplasm. (A-D) Representative paired sets from model training using nucleus (train (N=37) / test (N=5), mitochondria train (N=24) / test (N=5), and Cytoplasm train (N=61)/ test (N=9) models showing bright-field input, target signal and predicted signal (A), with train/validation rolling mean square error loss graphs for nucleus (B) mitochondria (C), and (D) Cytoplasm. (E) Correlation coefficient of target fluorescence (T) vs. predicted fluorescence (P) images paired with correlation coefficient from brightfield signal (S) for the organelles that were used to train the models.

The trained model was used to predict the coordinates of these structures in NCI-N87 cells (Fig. 3A-C), allowing quantification of 65 organelle features (Supplementary Table 2), including their volumes, areas and symmetry. We identified 11 of these features which differed significantly across cell cycle classes as defined by FUCCI across all images (Anova p-value *≤* 0.01; Figure 3D and Supplementary Figure 2), suggesting that changes in these structures during the cell cycle are quantifiable with label-free imaging.

**Figure 3.**
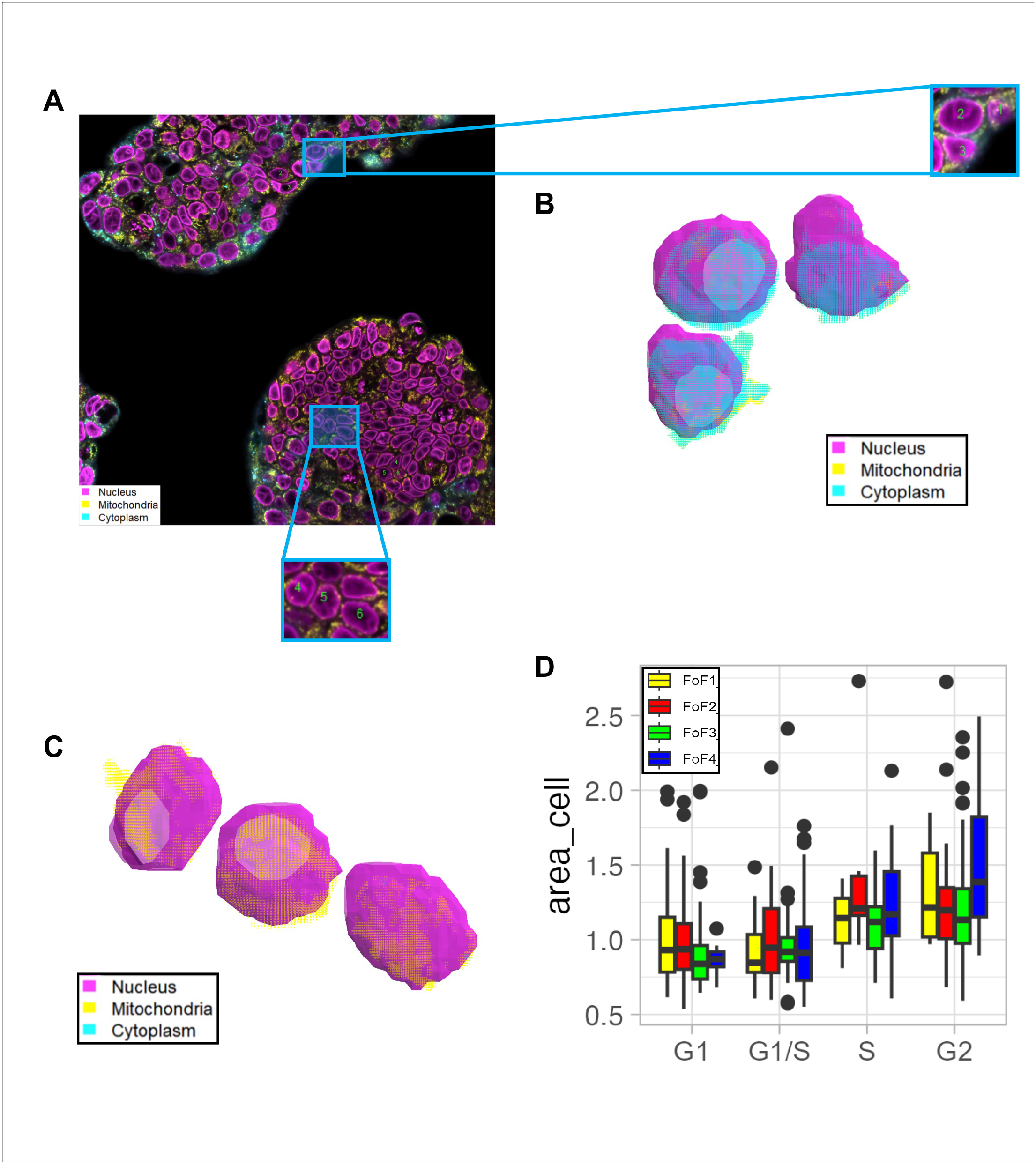
Label-free quantification of nucleus, cytoplasm and mitochondria at single cell resolution. A: Predicted nuclei, mitochondria and cytoplasm signals. B: Nuclei, mitochondria and cytoplasm of three representative cells from G1 or S phase. C: Nuclei, mitochondria and cytoplasm of three representative cells from G2/M phase. (B,C) Every data point is color-coded according to its organelle class. D: Cell features quantified from label free imaging correlate with cell cycle classes as defined by FUCCI. Color code indicates field of view (1-4). Data and code to produce this figure can be found on the project GitHub repository (Methods 4).

As a first test of this hypothesis, we used the FUCCI-based classification to train an SVM to predict discrete cell cycle state from the 11 nuclei-, mitochondria- and cytoplasm features (Supplementary Figure 2). The performance of the trained classifier was evaluated on an independent test set consisting of 373 unseen cells (Figure 4A). The classifier achieved an average accuracy *>* 0.7 across the four images for G1, G1/S transitional and G2M cells (Figure 4B). Classification accuracy was lower (median ≈ 0.6) for S-phase (Figure 4B).

**Figure 4.**
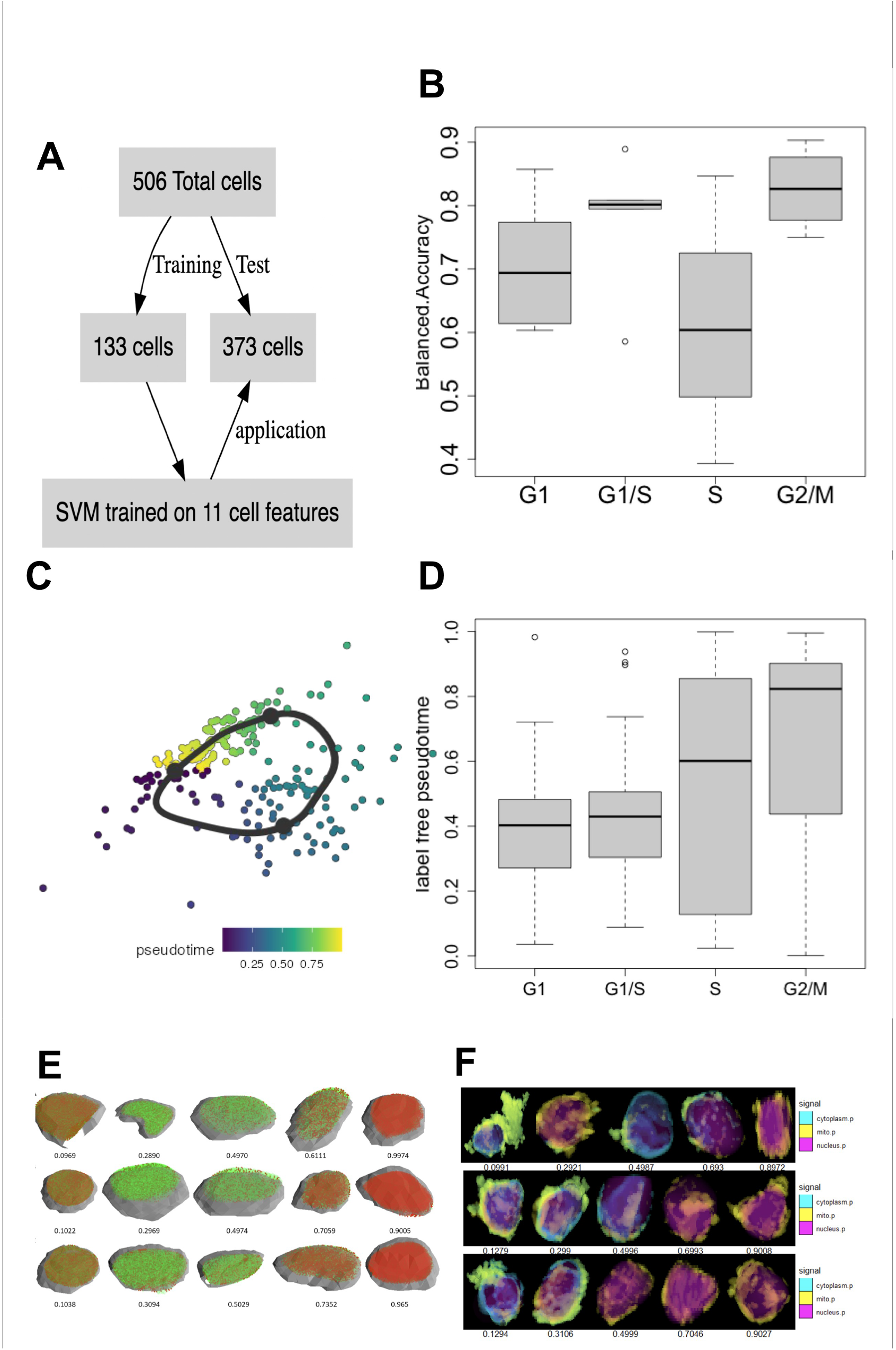
Characterization of discrete and continuous cell cycle progression in NCI-N87 with label free imaging. A: Supervised approach to predict discrete cell cycle state from nuclei, mitochondria and cytoplasm features. B: Performance of trained classifier on test set. C: Unsupervised approach to predict continuous cell cycle progression from nuclei, mitochondria and cytoplasm features. D: Pseudotime derived from label free imaging features is differentially distributed across the four FUCCI-informed cell cycle classes. E: representative cell across cell cycle: fucci. F: representative cell across cell cycle: label free. (E,F) x-axis indicates pseudotime. Data and code to produce this figure can be found on the project GitHub repository (Methods 4).

We also used the 11 cell organelle features as input to the Angle method for inferring pseudotime trajectories (Figure 4C). Similar to the FUCCI-derived pseudotime, the pseudotime inferred from label-free imaging was also correlated with and differentially distributed across the four discrete cell cycle classes (Figure 4D and Supplementary Figure 4).

Compared to FUCCI-derived pseudotime, pseudotime inferred from label-free imaging had a higher variance, possibly indicating a higher temporal resolution on the cell cycle. This approach allowed us to view virtual animations of the cell cycle by sampling cells representative across the entire pseudotime spectrum (Figure 4E,F).

### Cell cycle pseudotime links imaged to sequenced cells

Knowing a cell’s precise point on the cell cycle continuum is a novel opportunity to accomplish a challenging goal: integrating live-cell imaging with single-cell sequencing.

Well established and widely adopted methods for pseudotime inference from sequencing data exist and have been described in detail elsewhere, e.g. [29, 30, 31, 32]. These provide the opportunity to use the pseudotime derived herein from imaging in order to map imaged cells onto sequenced cells. To achieve this, we used single cell RNA sequencing (scRNAseq) data previously published for NCI-N87 [15]. A total of 1,076 genes involved in cell cycle, with highly variable expression among the 738 sequenced NCI-N87 cells, were prioritized for pseudotime inference with the Angle method (Figure 5A).

**Figure 5.**
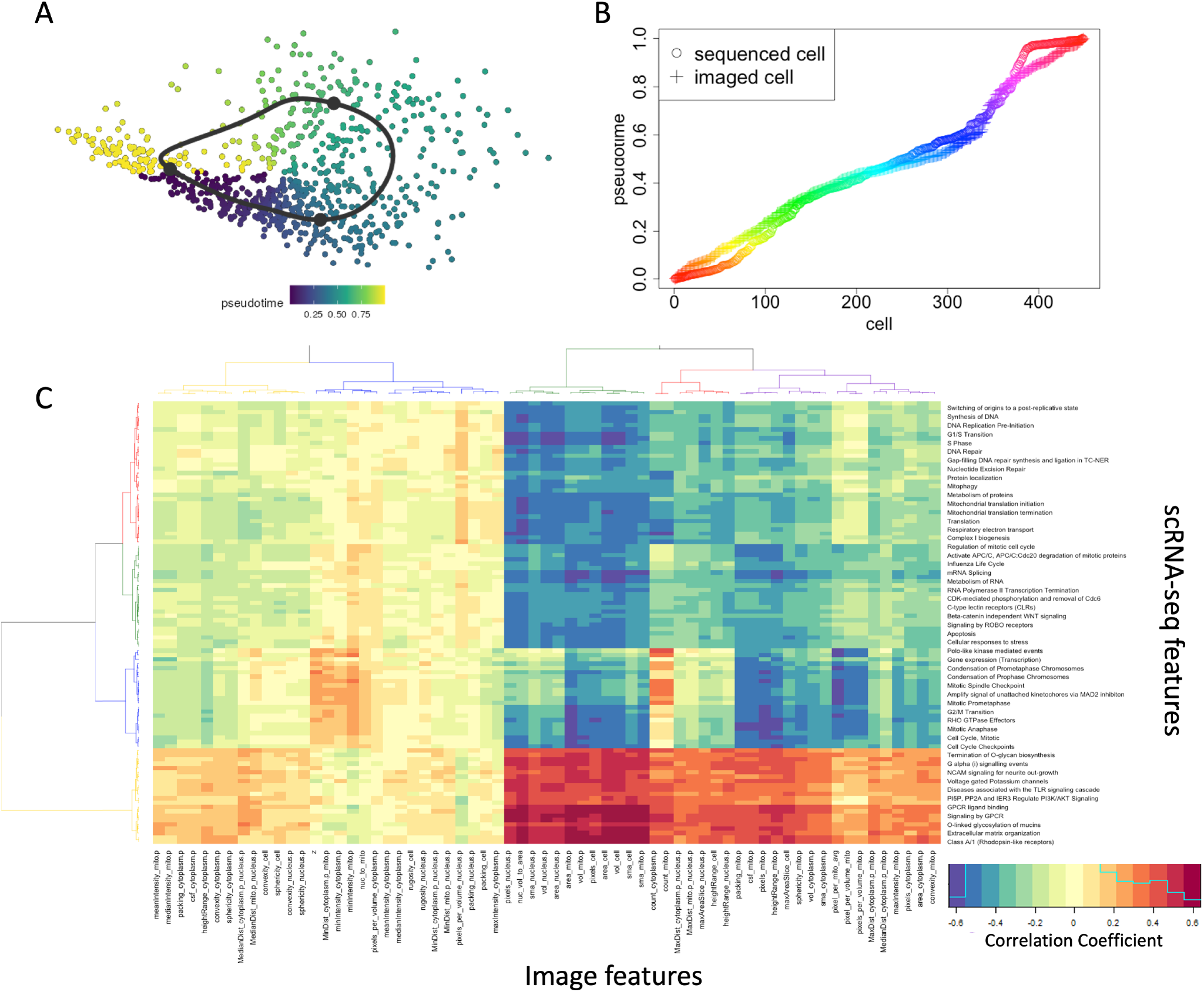
Sequencing imaging integration. A: Sequencing derived pseudotime of cell cycle progression. B: Co-clustering of sequenced and imaged cells based on pseudotime. Color code indicates cluster membership. C: Correlation between pathway activity and imaging-derived features. Data and code to produce this figure can be found on the project GitHub repository (Methods 4).

The distributions of sequencing- and imaging-derived pseudotimes were similar, and were co-clustered using DBSCAN (Figure 5B and Methods 4.10). This approach identified 225 clusters, 99% of which contained both, sequenced and imaged cell representatives (Figure 5B), further referred to as multi-type clusters. The average number of sequenced and imaged cells per cluster was 3.4 and 2.4 respectively. For each multi-type cluster we calculated the mean pathway activity among sequenced cell members and the mean organelle features among imaged cell members. Comparing all imaging- and sequencing derived feature pairs across clusters identified 4,484 significantly correlated feature pairs (Pearson r 0.38; Bonferroni-corrected p-value*<* 0.05; Figure 5C and Supplementary Table 1).

Of the top 150 pathways with the highest correlation coefficients (Supplementary Fig. 5), 24 were Signaling by GPCR, GPCR ligand binding, or GPCR downstream signaling; 18 were involved in G1, S or G1/S transition; and 18 were stages of mitosis. GPCR pathways had positive correlations with the area and volume of the cell, mitochondria and nucleus, as well as the nuclear volume to area ratio. G1/S pathways had negative correlations with the area and volume of the cell, mitochondria and the nuclear volume to area ratio. M pathways had a strong negative correlation with the area of mitochondria.

Overall, associations between pathway activities and cell morphology were evident in three large clusters. Pathways from the first cluster (indicated in yellow in Fig. 5C), are largely responsible for the formation of the extracellular matrix [33, 34]. These pathways also contribute to the organization of the ECM and collagen production. There are elements of cell growth and proliferation regulation, in part by downregulating the P13K/AKT pathway [35]. Cells which express these pathways tend to be large and round and have large volume and area of the nucleus, mitochondria, cytoplasm.

The second cluster included pathways that were almost entirely dedicated to mitosis (blue in Fig. 5C). They regulate the G2/M transition, each stage of M phase, the actin cytoskeleton components; the packaging, cohesion, and separation of sister chromatids; as well as other aspects of the mitotic spindle and chromosome organization. Cells which have high expression of these pathways will have slightly higher cytoplasmic and mitochondrial intensities, lower mitochondrial resolution, and elevated mitochondrial counts. Indeed, the number of mitochondria has been described to increase during G2 and M phases compared to G1/S due to the fusion of mitochondria in G1/S and subsequent fission in G2/M [36].

The third cluster (indicated in red and green in Fig. 5C), consisted of pathways that play a major role in cell cycle transitioning and replication of genetic material. A selection of these elements includes the conversion from S to M by Orc1 removal [37] and DNA synthesis, replication, damage recognition, and repair; mRNA processing, apoptosis, environmental responsiveness, angiogenesis and motility, and cell cycle protein regulation. Cells expressing these pathways were smaller, had a low maximum intensity of mitochondria, low area of cytoplasm, low convexity of mitochondria, and substantially lower volume and area of the mitochondria and nucleus.

Taken together these associations between imaging- and transcriptome derived features pairs are in line with decades of research unraveling how the transcriptome influences cell shape [11, 9, 10], scaling [7, 12], compartmentalization [6, 5, 8] and protein localization [38].

## 3 Discussion

A combination of unsupervised and supervised classification methods have been developed to classify imaged cells to one of three possible cell cycle states (G1, S or G2M). Some studies however suggest that gene expression signatures of cell state transitions occur as a continuous process, rather than in abrupt steps [30, 39, 40]. We have shown that both, 3D imaging of FUCCI cells as well as a high resolution on the 3D subcellular architecture of cells can provide a quantitative description of cell cycle progression, allowing us to move beyond the classification of cells into discrete states. While it is unclear which of the two – FUCCI- or organelle features – provide a higher temporal resolution on cell cycle progression, it is noteworthy that quantification of FUCCI features per nucleus were dependent on the accurate segmentation of nuclei, which in turn were derived from semantic segmentation of label-free images. Ultimately, it is reasonable to hypothesize that the highest temporal resolution can be achieved by combining both methodologies. Verifying this hypothesis will require applying the approach presented herein to a live cell imaging experiment spanning multiple days.

In contrast to imaging data, for which methods for classification of cell cycle state are scarce [41, 42], several methods for cell cycle inference from sequencing data exist [29, 30, 31, 43] and are widely adopted. We have for the first time integrated sequencing and imaging derived cell cycle pseudotimes for mapping clusters of imaged cells to sequenced cell clusters from the same gastric cancer cell line. For this experiment, sequenced and imaged cells were obtained from different timepoints. The next step will be to repeat this approach within a longer-term live cell imaging experiments, wherein cells are sampled intermittently for sequencing.

Knowing the entire transcriptome of a cell will always require killing the cell. The ability to assign the transcriptome state of a lysed cell to its closest living relative (which is still actively growing and expanding), would be unprecedented and would open the door for genotype-phenotype mapping at single cell resolution, forward in time. The growing field of spatial transcriptomics [44], while simplifying mapping between imaged cells and their transcriptomes, cannot accomplish this particular task because it requires killing all spatially adjacent cells for sequencing. This means that spatial transcriptomics cannot be used for learning to predict phenotypes forward in time, but could only be leveraged for retrospective phenotypic interpretation. The phenotypic interpretation of genomes and transcriptomes is the bottleneck to progress in medicine, it lags far behind manipulation and quantification. By understanding how the transcriptomes and genomes of co-existing cells diverge, how these divergent populations compete or cooperate, one can learn to predict the long-term consequence of exposing them to different therapeutic environments.

## 4 Methods

Our proposed approach for inferring cell’s position along the cell cycle continuum from 3D images is depicted in Figure 6. In the following subsections, we illustrate each component of the proposed approach. Publicly available code and data is located at https://github.com/MathOnco/CellCycle4DataIntegration

**Figure 6.**
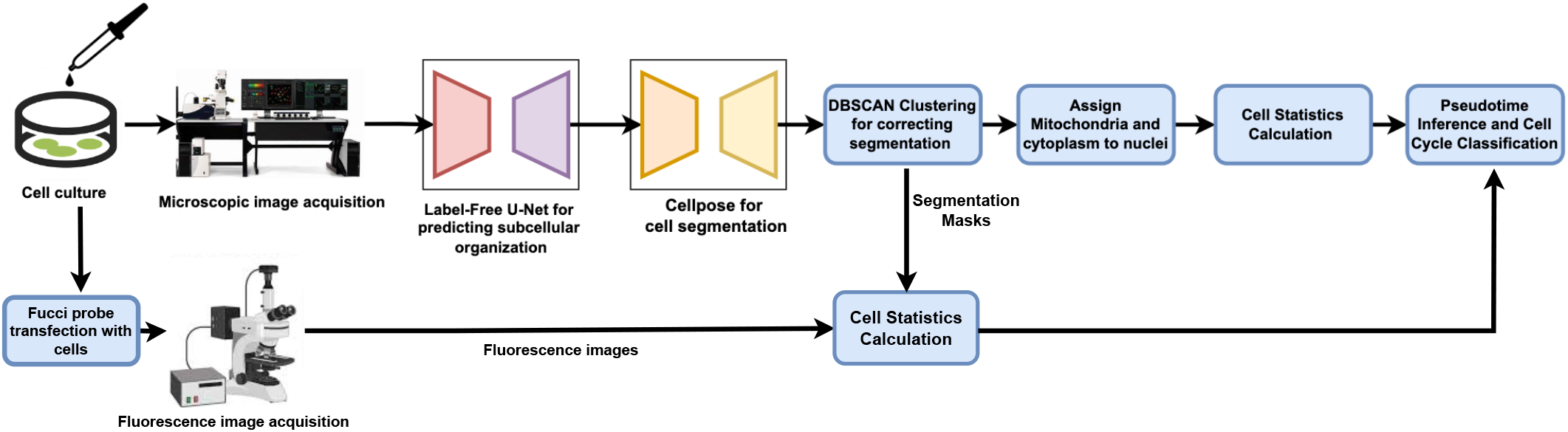
An illustration of our proposed approach for cell cycle classification and assigning sequenced to imaged cells.

### 4.1 Transfection of NCI-N87 cells with FUCCI-vector plasmids

For cell cycle-phase visualization, lentiviral FUCCI (fluorescent ubiquitination-based cell cycle indicator) expression system was used. The PIP Fucci vector (Addgene, Plasmid #118616) encoding the FUCCI probe was co-transfected with the packaging plasmids into HEK 293T cells. Supernatant from the culture medium containing high-titer viral solutions were collected and used for transduction into NCI-N87 cells. PIP Fucci labels G1 cells with mVenus (green) and S-G2-M cells with (mCherry). Cells with stable integration of the plasmid were established by FACS with both green and red channels.

#### Cell culture

NCI-N87 were seeded in µ-Slide 8 Well, ibiTreat - Tissue Culture Treated Polymer Coverslip (Fisher Scientific) using 5 ×10^4^ or 1 × 10^5^ cells in 300ul of RPMI-1640 with 10% FBS and 1% penicillin streptomycin. Cells were treated with 0.3ul or 0.6ul BioTracker 488 Nuclear per 300ul for 3hrs or 70-120nM BioTracker 405 Mito for 6hrs (Millipore Sigma). Cells were washed 3x with warm PBS and resuspended in complete growth media for imaging.

### 4.2 Microscopy

A confocal microscope (Leica TCS SP8) equipped with a 63X/1.4-NA oil-immersion objective (Leica Apochromat × 100/1.4 W) was used for image acquisition. The 3D cell images were recorded in LAS X 3.5.7 using Photomultiplier Tube detectors, resulting in a pixel size of 0.232 *µ*m and Z-interval of 0.29 *µ*m. We collected 70 z slices of target fluorescence dye (cytoplasm, mitochondria or nuclei) and brightfield signal using a 400 Hz scan speed. For each field of view, we imaged an area of 56,644 *µ*m^2^, which took approximately 3 min for the brightfield and the fluorescence channels.

### 4.3 Training and application of a U-Net convolutional neural network to predict subcellular organization

Stacks of 70 brightfield and fluorescence 16-bit images were processed into “.ome.tif” files containing NumPy arrays. We trained a previously developed label-free U-Net convolutional neural network [16], on these 3D images containing the nuclei or mitochondria of NCI-N87 to calculate the spatial coordinates of nucleus and mitochondria in each imaged cell. All models were trained using number of batches=8 for 3D patches of 128 × 128 × 32p^3^(XYZ), the Adam optimizer with a learning rate of 0.001 and with beta, *β*_1_ and, *β*_2_ of 0.9 and 0.999 respectively for 150,000 minibatch iterations. The model training pipeline was implemented in Python using PyTorch on a Nvidia DGX A100 Tesla V100.

The trained model was applied to 3D brightfield live-cell imaging of NCI-N87 cells for nine hours at a three hour interval, acquiring a total of four images. Image acquisition was performed as described above (Methods 4.2). The nuclei and mitochondria predicted by the trained model were segmented using Cellpose [27] – a generalist model that segments cells from a wide range of image types including 2D and 3D images. Cellpose is based on a U-Net architecture with residual blocks. In 3D cell segmentation, testing of the fine-tuned pretrained Cellpose model (i.e., cytotorch 2) was completed where a gradient was generated for xy, yz, xz slices, independently. Then the gradient was averaged to obtain the final prediction. This approach allows testing 3D images on a 2D based deep learning model.

### 4.4 Correcting Nuclei Segmentation

We correct segmentation by merging IDs belonging to the same cell using 3D imaging data. We read the center coordinates of cells from a CSV file and scale the z-coordinates according to the specified Z-stack distance. Using DBSCAN clustering on the x, y, and z coordinates, we identify clusters of points representing individual cells, filtering out noise.

For each identified cell cluster, the function gathers coordinates associated with the new cell from the original segmentation files. If a cell does not have sufficient z-stack representation, it is excluded. The function then calculates the centroids of the newly identified cells.

### 4.5 Assigning mitochondria and cytoplasm to nuclei

We assign mitochondrial and cytoplasmic compartments to nuclei based on 3D imaging data. For each organelle (mitochondria, cytoplasm), the function reads the corresponding TIF image, identifies pixels with intensity above the 90th percentile, and records their coordinates and signal values. Nucleus coordinate files are then loaded and associated with their respective cells.

For each nucleus, we expand the bounding box around it, identify organelle pixels within this expanded region, and assign these pixels to the nucleus. DBSCAN clustering is performed on the combined coordinates (nucleus, mitochondria, and cytoplasm) for each cell. The cluster containing the nucleus is identified and organelle pixels in this cluster are assigned to the corresponding cell. We correct for doubly assigned coordinates by removing ambiguous assignments. This method ensures accurate spatial assignment of mitochondrial and cytoplasmic compartments to individual nuclei for further analysis.

### 4.6 Cell Statistics Calculation

Statistics are calculated for each cell based on its segmented coordinates and signal intensities. Statistics related to nucleus shape and size are computed, including area, volume, fractal dimension, rugosity, and height range. Next, organelle-specific statistics are computed. For each signal type (e.g., mitochondria, cytoplasm), we iterate through unique organelles within the cell and computes statistics related to shape, size, and spatial relationships, including area, volume, convexity, packing, sphericity, and distance to other organelles, as well as intensity-based features (mean, median, maximum, and minimum intensity values). R packages ‘geometry’, ‘habtools’ and ‘misc3d’ are used to calculate these statistics.

Additional statistics calculated include average pixels per mitochondria, pixel density per volume for each organelle, and ratios between organelle volumes.

### 4.7 Pseudotime Inference

Cell features derived from either i) FUCCI or from ii) label-free imaging are used on multiple fields of view to infer pseudotime. For both i) and ii), each feature vector is divided by its median across all cells and log-transformed prior to inference. Trajectory inference is conducted using the Angle method, while employing principal component analysis (PCA) for dimensionality reduction. To evaluate potential batch effects, pseudotime inference is performed on all fields of view combined as well as on each field of view separately.

### 4.8 Taining an SVM to classify cells to cell cycle phases

We perform cell cycle phase classification using Support Vector Machine (SVM) and evaluate the model’s performance as follows. An SVM with a radial basis kernel is trained using cell features derived from label free imaging as input, and the four cell cycle classes (inferred from FUCCI imaging) as labels. This trained SVM model is subsequently used to predict cell cycle phases on test data from different cells across all available fields of view.

### 4.9 Quantification of pathway activity from gene expression

Gene set variation analysis (GSVA) is performed to compute the activities of 1,119 pathways from the REACTOME database [45], based on single-cell RNA sequencing (scRNA-seq) of 738 NCI-N87 cells. R-function “gsva” was used from the R-package GSVA.

### 4.10 Assigning sequenced to imaged cells based on pseudotime

This segment aims to co-cluster image and sequencing statistics for further analysis. Pseudotimes inferred from sequencing (P-seq) and from imaging (P-img) are combined into a one-dimensional vector and a density-based clustering algorithm (DBSCAN) is applied to the combined vector (R-function ‘db-scan::dbscan’ is used with *eps* = 0.001 and *minPts* = 2). Clusters that consist solely of one data type (either “P-seq” or “P-img”) and noise are filtered out.

For each remaining cluster we calculate two averages: the average pathway activity profile across all sequenced cell members of that cluster and the average organelle feature profile across all imaged cell members of that cluster. Subsequently, Pearson correlation coefficients between all possible feature-pairs are calculated.

## Acknowledgements

We thank Ana Gomes and Stanislav Drapela for providing the FUCCI-vector plasmids; and Florian Karreth for advice on the cell organelle dyes used in this study.

## References

[1] U. Ben-David, B. Siranosian, G. Ha, H. Tang, Y. Oren, K. Hinohara, C. A. Strathdee, J. Dempster, N. J. Lyons, R. Burns, A. Nag, G. Kugener, B. Cimini, P. Tsvetkov, Y. E. Maruvka, R. O’Rourke, A. Garrity, A. A. Tubelli, P. Bandopadhayay, A. Tsherniak, F. Vazquez, B. Wong, C. Birger, M. Ghandi, A. R. Thorner, J. A. Bittker, M. Meyerson, G. Getz, R. Beroukhim, and T. R. Golub. Genetic and transcriptional evolution alters cancer cell line drug response. 560(7718):325–330. PMID: 30089904.

[2] U. Ben-David, G. Ha, Y.-Y. Tseng, N. F. Greenwald, C. Oh, J. Shih, J. M. McFarland, B. Wong, J. S. Boehm, R. Beroukhim, and T. R. Golub. Patient-derived xenografts undergo murine-specific tumor evolution. 49(11):1567–1575. PMID: 28991255.

[3] N. Andor, T. A. Graham, M. Jansen, L. C. Xia, C. A. Aktipis, C. Petritsch, H. P. Ji, and C. C. Maley. Pan-cancer analysis of the extent and consequences of intratumor heterogeneity. 22(1):105–113. PMID: 26618723.

[4] M. Chen, B. Zhang, W. Topatana, J. Cao, H. Zhu, S. Juengpanich, Q. Mao, H. Yu, and X. Cai. Classification and mutation prediction based on histopathology H&E images in liver cancer using deep learning. 4(1):1–7.

[5] C.-Y. Wu, P. A. Rolfe, D. K. Gioffrd, and G. R. Fink. Control of Transcription by Cell Size. 8(11):e1000523.

[6] D. Bausch-Fluck, U. Goldmann, S. Müller, M. van Oostrum, M. Müller, O. T. Schubert, and B. Wollscheid. The in silico human surfaceome. 115(46):E10988–E10997. PMID: 30373828.

[7] Y.-H. M. Chan and W. F. Marshall. Scaling properties of cell and organelle size. 6(2):88–96. PMID: 20885855.

[8] Y.-H. M. Chan and W. F. Marshall. How cells know the size of their organelles. 337(6099):1186–1189. PMID: 22955827.

[9] I. Nassiri and M. N. McCall. Systematic exploration of cell morphological phenotypes associated with a transcriptomic query. 46(19):e116. PMID: 30011038.

[10] P. Rangamani, A. Lipshtat, E. U. Azeloglu, R. C. Calizo, M. Hu, S. Ghassemi, J. Hone, S. Scarlata, S. R. Neves, and R. Iyengar. Decoding information in cell shape. 154(6):1356–1369. PMID: 24034255.

[11] M. F. A. Cutiongco, B. S. Jensen, P. M. Reynolds, and N. Gadegaard. Predicting gene expression using morphological cell responses to nanotopography. 11(1):1384.

[12] R. J. Kimmerling, S. M. Prakadan, A. J. Gupta, N. L. Calistri, M. M. Stevens, S. Olcum, N. Cermak, R. S. Drake, K. Pelton, F. De Smet, K. L. Ligon, A. K. Shalek, and S. R. Manalis. Linking single-cell measurements of mass, growth rate, and gene expression. 19(1):207. PMID: 30482222.

[13] K. Lane, D. Van Valen, M. M. DeFelice, D. N. Macklin, T. Kudo, A. Jaimovich, A. Carr, T. Meyer, D. Pe’er, S. C. Boutet, and M. W. Covert. Measuring Signaling and RNA-Seq in the Same Cell Links Gene Expression to Dynamic Patterns of NF-KB Activation. 4(4):458–469.e5. PMID: 28396000.

[14] J. Yuan, J. Sheng, and P. A. Sims. SCOPE-Seq: a scalable technology for linking live cell imaging and single-cell RNA sequencing. 19(1):227. PMID: 30583733.

[15] N. Andor, B. T. Lau, C. Catalanotti, A. Sathe, M. Kubit, J. Chen, C. Blaj, A. Cherry, C. D. Bangs, S. M. Grimes, C. J. Suarez, and H. P. Ji. Joint single cell DNA-seq and RNA-seq of gastric cancer cell lines reveals rules of in vitro evolution. 2(2). PMID: 32215369.

[16] C. Ounkomol, S. Seshamani, M. M. Maleckar, F. Collman, and G. R. Johnson. Label-free prediction of three-dimensional fluorescence images from transmitted-light microscopy. 15(11):917–920.

[17] B. Alberts, A. Johnson, J. Lewis, M. Raff, K. Roberts, and P. Walter. The Compartmentalization of Cells.

[18] H. X. Chao, C. E. Poovey, A. A. Privette, G. D. Grant, H. Y. Chao, J. G. Cook, and J. E. Purvis. Orchestration of DNA Damage Checkpoint Dynamics across the Human Cell Cycle. 5(5):445–459.e5. PMID: 29102360.

[19] C. A. Yates, M. J. Ford, and R. L. Mort. A Multi-stage Representation of Cell Proliferation as a Markov Process. 79(12):2905–2928. PMID: 29030804.

[20] H. X. Chao, R. I. Fakhreddin, H. K. Shimerov, K. M. Kedziora, R. J. Kumar, J. Perez, J. C. Limas, G. D. Grant, J. G. Cook, G. P. Gupta, and J. E. Purvis. Evidence that the human cell cycle is a series of uncoupled, memoryless phases. 15(3):e8604. PMID: 30886052.

[21] K. R. Ghusinga, C. A. Vargas-Garcia, and A. Singh. A mechanistic stochastic framework for regulating bacterial cell division. 6:30229. PMID: 27456660.

[22] C. Garmendia-Torres, O. Tassy, A. Matifas, N. Molina, and G. Charvin. Multiple inputs ensure yeast cell size homeostasis during cell cycle progression. 7:e34025.

[23] K. E. Coleman, G. D. Grant, R. A. Haggerty, K. Brantley, E. Shibata, B. D. Workman, A. Dutta, D. Varma, J. E. Purvis, and J. G. Cook. Sequential replication-coupled destruction at G1/S ensures genome stability. 29(16):1734–1746. PMID: 26272819.

[24] W. Saelens, R. Cannoodt, H. Todorov, and Y. Saeys. A comparison of single-cell trajectory inference methods. 37(5):547–554. PMID: 30936559.

[25] J. K. Tung, K. Berglund, C.-A. Gutekunst, U. Hochgeschwender, and R. E. Gross. Bioluminescence imaging in live cells and animals. 3(2):025001. PMID: 27226972.

[26] O. Ronneberger, P. Fischer, and T. Brox. U-Net: Convolutional Networks for Biomedical Image Segmentation.

[27] C. Stringer, T. Wang, M. Michaelos, and M. Pachitariu. Cellpose: a generalist algorithm for cellular segmentation. 18(1):100–106.

[28] M. Hahsler, M. Piekenbrock, and D. Doran. dbscan: Fast Density-Based Clustering with R. 91:1–30.

[29] K. Street, D. Risso, R. B. Fletcher, D. Das, J. Ngai, N. Yosef, E. Purdom, and S. Dudoit. Slingshot: cell lineage and pseudotime inference for single-cell transcriptomics. 19(1):477.

[30] C. J. Hsiao, P. Tung, J. D. Blischak, J. E. Burnett, K. Barr, K. K. Dey, M. Stephens, and Y. Gilad. Characterizing and inferring quantitative cell cycle phase in single-cell RNA-seq data analysis. page 526848.

[31] N. Leng, L.-F. Chu, C. Barry, Y. Li, J. Choi, X. Li, P. Jiang, R. M. Stewart, J. A. Thomson, and C. Kendziorski. Oscope identifies oscillatory genes in unsynchronized single-cell RNA-seq experiments. 12(10):947–950.

[32] M. J. Povinelli and J. A. Robinson. Integrating Design Thinking into an Experiential Learning Course for Freshman Engineering Students.

[33] A. Varki, R. Cummings, J. Esko, H. Freeze, G. Hart, and J. Marth. O-Glycans. In Essentials of Glycobiology. Cold Spring Harbor Laboratory Press.

[34] J. Casale and J. S. Crane. Biochemistry, Glycosaminoglycans. In StatPearls. StatPearls Publishing.

[35] A. C. Lloyd. The Regulation of Cell Size. 154(6):1194–1205.

[36] P. Mishra and D. C. Chan. Mitochondrial dynamics and inheritance during cell division, development and disease. 15(10):634–646. PMID: 25237825.

[37] C.-J. Li and M. L. DePamphilis. Mammalian Orc1 Protein Is Selectively Released from Chromatin and Ubiquitinated during the S-to-M Transition in the Cell Division Cycle. 22(1):105–116. PMID: 11739726.

[38] A. Mondal and J. Hu. Network based prediction of protein localisation using diffusion kernel. 9(4):386– 400. PMID: 25757246.

[39] S. Pauklin and L. Vallier. The cell-cycle state of stem cells determines cell fate propensity. 155(1):135– 147. PMID: 24074866.

[40] N. T. Ingolia and A. W. Murray. The ups and downs of modeling the cell cycle. 14(18):R771–777. PMID: 15380091.

[41] X. Jin, Y. Zou, and Z. Huang. An Imbalanced Image Classification Method for the Cell Cycle Phase. 12(6):249.

[42] Y. Nagao, M. Sakamoto, T. Chinen, Y. Okada, and D. Takao. Robust classification of cell cycle phase and biological feature extraction by image-based deep learning. 31(13):1346–1354.

[43] B. Povinelli, Q. Wills, N. Barkas, C. Booth, K. Campbell, A. Rodriguez-Meira, S. E. Jacobsen, C. Yau, and A. Mead. Integrated Single Cell Analysis Reveals Cell Cycle and Ontogeny Related Transcriptional Heterogeneity in Hscs. 64:S95–S96.

[44] A. Rao, D. Barkley, G. S. FranÇa, and I. Yanai. Exploring tissue architecture using spatial transcriptomics. 596(7871):211–220.

[45] B. Jassal, L. Matthews, G. Viteri, C. Gong, P. Lorente, A. Fabregat, K. Sidiropoulos, J. Cook, M. Gillespie, R. Haw, F. Loney, B. May, M. Milacic, K. Rothfels, C. Sevilla, V. Shamovsky, S. Shorser, T. Varusai, J. Weiser, G. Wu, L. Stein, H. Hermjakob, and P. D’Eustachio. The reactome pathway knowledgebase. 48:D498–D503. PMID: 31691815.

[46] S. Prabhakaran, C. Gatenbee, M. Robertson-Tessi, J. West, A. A. Beg, J. Gray, S. Antonia, R. A. Gatenby, and A. R. A. Anderson. Mistic: An open-source multiplexed image t-SNE viewer. 3(7).

