## Supplemental figures and tables for "Cell identity revealed by precise cell cycle state mapping links data modalities"

### Supplementary Information

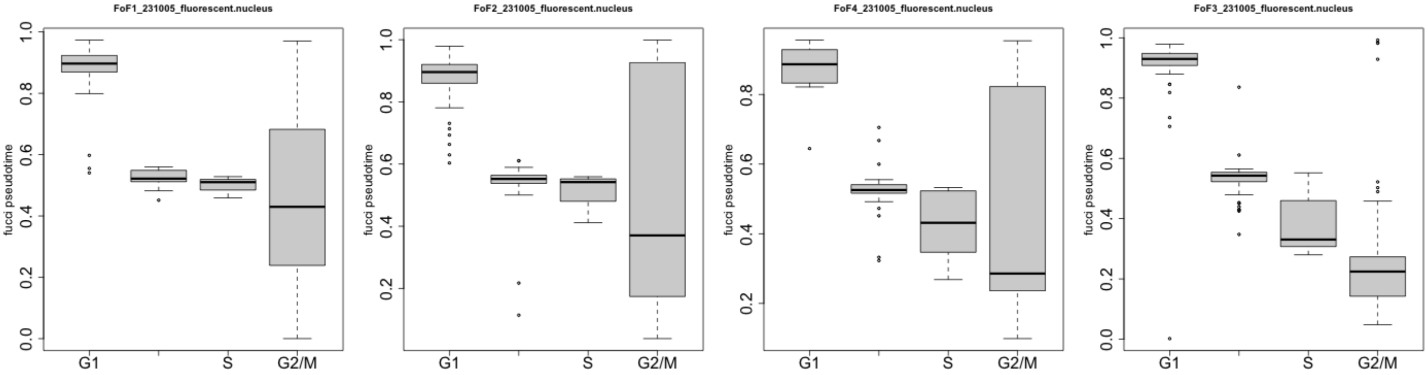


**Supplementary Figure 1:** Fucci-derived pseudtime is differentially distributed across the four cell cycle phases. .

| Pathway | Cell.Feature | r | pval |
|---|---|---|---|
| S Phase | nuc_vol_to_area | -0.64 | 0.00 |
| mRNA Splicing | area_mito.p | -0.62 | 0.00 |
| Resolution of Sister Chromatid Cohesion | pixel_per_mito_avg | -0.62 | 0.00 |
| G alpha (q) signalling events | area_cell | 0.62 | 0.00 |
| Processing of Capped Intron-Containing P... | area_mito.p | -0.62 | 0.00 |
| GPCR ligand binding | area_cell | 0.62 | 0.00 |
| Mitotic G1-G1/S phases | nuc_vol_to_area | -0.62 | 0.00 |
| Amplification of signal from unattached... | pixel_per_mito_avg | -0.61 | 0.00 |
| Amplification of signal from the kinetoc... | pixel_per_mito_avg | -0.61 | 0.00 |
| G alpha (q) signalling events | vol_cell | 0.61 | 0.00 |
| Signaling by GPCR | area_cell | 0.61 | 0.00 |
| Mitotic Spindle Checkpoint | pixel_per_mito_avg | -0.61 | 0.00 |
| GPCR ligand binding | vol_cell | 0.61 | 0.00 |
| G1/S Transition | nuc_vol_to_area | -0.61 | 0.00 |
| mRNA Splicing | area_cell | -0.61 | 0.00 |
| Signaling by GPCR | vol_cell | 0.60 | 0.00 |
| Cell Cycle, Mitotic | area_mito.p | -0.60 | 0.00 |
| RHO GTPase Effectors | pixels_mito.p | -0.60 | 0.00 |
| Processing of Capped Intron-Containing P... | area_cell | -0.60 | 0.00 |
| GPCR ligand binding | sma_cell | 0.60 | 0.00 |
| GPCR ligand binding | area_mito.p | 0.60 | 0.00 |
| Signaling by GPCR | sma_cell | 0.60 | 0.00 |
| Extracellular matrix organization | area_cell | 0.60 | 0.00 |
| G alpha (q) signalling events | sma_cell | 0.60 | 0.00 |
| mRNA Splicing | vol_cell | -0.60 | 0.00 |
| Signaling by GPCR | nuc_vol_to_area | 0.60 | 0.00 |
| Mitotic Prometaphase | pixel_per_mito_avg | -0.60 | 0.00 |
| GPCR ligand binding | sma_mito.p | 0.60 | 0.00 |
| The citric acid (TCA) cycle and respirat... | nuc_vol_to_area | -0.60 | 0.00 |
| Class A/1 (Rhodopsin-like receptors) | area_cell | 0.60 | 0.00 |
| Peptide ligand-binding receptors | area_cell | 0.60 | 0.00 |

| | | | |
|---|---|---|---|
| S Phase | vol_nucleus.p | -0.59 | 0.00 |
| Processing of Capped Intron-Containing P... | vol_cell | -0.59 | 0.00 |
| Cell Cycle | area_mito.p | -0.59 | 0.00 |
| Cell Cycle Checkpoints | area_mito.p | -0.59 | 0.00 |
| GPCR downstream signalling | area_cell | 0.59 | 0.00 |
| Mitotic Anaphase | pixels_mito.p | -0.59 | 0.00 |
| Mitotic Metaphase and Anaphase | pixels_mito.p | -0.59 | 0.00 |
| RHO GTPase Effectors | area_mito.p | -0.59 | 0.00 |
| S Phase | pixels_nucleus.p | -0.59 | 0.00 |
| O-linked glycosylation | area_cell | 0.59 | 0.00 |
| Class A/1 (Rhodopsin-like receptors) | vol_cell | 0.59 | 0.00 |
| Peptide ligand-binding receptors | vol_cell | 0.59 | 0.00 |
| GPCR downstream signalling | vol_cell | 0.59 | 0.00 |
| GPCR downstream signalling | sma_cell | 0.59 | 0.00 |
| Extracellular matrix organization | vol_cell | 0.59 | 0.00 |
| GPCR ligand binding | nuc_vol_to_area | 0.59 | 0.00 |
| Synthesis of DNA | nuc_vol_to_area | -0.59 | 0.00 |
| G alpha (q) signalling events | nuc_vol_to_area | 0.59 | 0.00 |
| GPCR downstream signalling | nuc_vol_to_area | 0.59 | 0.00 |
| Separation of Sister Chromatids | pixels_mito.p | -0.59 | 0.00 |
| mRNA Splicing | vol_mito.p | -0.59 | 0.00 |
| RHO GTPases Activate Formins | pixel_per_mito_avg | -0.58 | 0.00 |
| O-linked glycosylation | area_mito.p | 0.58 | 0.00 |
| O-linked glycosylation | nuc_vol_to_area | 0.58 | 0.00 |
| S Phase | area_nucleus.p | -0.58 | 0.00 |
| M Phase | pixels_mito.p | -0.58 | 0.00 |
| GPCR ligand binding | vol_mito.p | 0.58 | 0.00 |
| Cyclin A:Cdk2-associated events at S pha... | nuc_vol_to_area | -0.58 | 0.00 |
| O-linked glycosylation | vol_cell | 0.58 | 0.00 |
| DNA Replication | nuc_vol_to_area | -0.58 | 0.00 |
| Signaling by GPCR | pixels_nucleus.p | 0.58 | 0.00 |
| G alpha (q) signalling events | sma_mito.p | 0.58 | 0.00 |
| Extracellular matrix organization | area_mito.p | 0.58 | 0.00 |
| Processing of Capped Intron-Containing P... | vol_mito.p | -0.58 | 0.00 |
| O-linked glycosylation | sma_cell | 0.58 | 0.00 |
| Cyclin A/B1/B2 associated events during ... | pixel_per_mito_avg | -0.58 | 0.00 |
| mRNA Splicing | sma_mito.p | -0.58 | 0.00 |
| Mitotic G1-G1/S phases | pixels_nucleus.p | -0.58 | 0.00 |
| Metabolism of RNA | area_cell | -0.58 | 0.00 |
| Metabolism of RNA | vol_cell | -0.58 | 0.00 |
| Mitotic G1-G1/S phases | vol_cell | -0.58 | 0.00 |
| Mitotic Anaphase | area_mito.p | -0.57 | 0.00 |
| G alpha (q) signalling events | pixels_nucleus.p | 0.57 | 0.00 |
| Mitotic Metaphase and Anaphase | area_mito.p | -0.57 | 0.00 |
| Signaling by GPCR | sma_mito.p | 0.57 | 0.00 |
| Class A/1 (Rhodopsin-like receptors) | nuc_vol_to_area | 0.57 | 0.00 |
| Peptide ligand-binding receptors | nuc_vol_to_area | 0.57 | 0.00 |
| Signaling by GPCR | area_mito.p | 0.57 | 0.00 |
| RHO GTPase Effectors | pixels_cell | -0.57 | 0.00 |
| GPCR downstream signalling | pixels_nucleus.p | 0.57 | 0.00 |

| | | | |
|---|---|---|---|
| G alpha (q) signalling events | area_mito.p | 0.57 | 0.00 |
| Mitotic G1-G1/S phases | area_cell | -0.57 | 0.00 |
| The citric acid (TCA) cycle and respirat... | pixels_nucleus.p | -0.57 | 0.00 |
| mRNA Splicing | heightRange_mito.p | -0.57 | 0.00 |
| Class A/1 (Rhodopsin-like receptors) | area_mito.p | 0.57 | 0.00 |
| Peptide ligand-binding receptors | area_mito.p | 0.57 | 0.00 |
| S Phase | vol_cell | -0.57 | 0.00 |
| M Phase | area_mito.p | -0.57 | 0.00 |
| DNA Repair | nuc_vol_to_area | -0.57 | 0.00 |
| Class A/1 (Rhodopsin-like receptors) | vol_mito.p | 0.57 | 0.00 |
| Peptide ligand-binding receptors | vol_mito.p | 0.57 | 0.00 |
| Mitotic G1-G1/S phases | vol_nucleus.p | -0.57 | 0.00 |
| G alpha (q) signalling events | vol_mito.p | 0.57 | 0.00 |
| Class A/1 (Rhodopsin-like receptors) | pixels_cell | 0.57 | 0.00 |
| Peptide ligand-binding receptors | pixels_cell | 0.57 | 0.00 |
| Mitochondrial translation elongation | nuc_vol_to_area | -0.57 | 0.00 |
| Separation of Sister Chromatids | area_mito.p | -0.57 | 0.00 |
| Translation | nuc_vol_to_area | -0.57 | 0.00 |
| G1/S Transition | pixels_nucleus.p | -0.57 | 0.00 |
| O-linked glycosylation | sma_mito.p | 0.57 | 0.00 |
| S Phase | area_cell | -0.57 | 0.00 |
| Extracellular matrix organization | sma_cell | 0.57 | 0.00 |
| G1/S Transition | vol_nucleus.p | -0.57 | 0.00 |
| Polo-like kinase mediated events | pixel_per_mito_avg | -0.56 | 0.00 |
| Signaling by Rho GTPases | pixels_mito.p | -0.56 | 0.00 |
| Mitotic Anaphase | heightRange_mito.p | -0.56 | 0.00 |
| Processing of Capped Intron-Containing P... | heightRange_mito.p | -0.56 | 0.00 |
| Mitochondrial protein import | nuc_vol_to_area | -0.56 | 0.00 |
| Signaling by GPCR | vol_nucleus.p | 0.56 | 0.00 |
| G2/M Transition | pixels_mito.p | -0.56 | 0.00 |
| Extracellular matrix organization | sma_mito.p | 0.56 | 0.00 |
| Mitotic Metaphase and Anaphase | heightRange_mito.p | -0.56 | 0.00 |
| GPCR downstream signalling | sma_mito.p | 0.56 | 0.00 |
| G1/S Transition | vol_cell | -0.56 | 0.00 |
| M Phase | pixel_per_mito_avg | -0.56 | 0.00 |
| Cell Cycle Checkpoints | heightRange_mito.p | -0.56 | 0.00 |
| Cell Cycle, Mitotic | vol_mito.p | -0.56 | 0.00 |
| Metabolism of RNA | area_mito.p | -0.56 | 0.00 |
| Respiratory electron transport | nuc_vol_to_area | -0.56 | 0.00 |
| Mitochondrial translation termination | nuc_vol_to_area | -0.56 | 0.00 |
| Respiratory electron transport, ATP synt... | nuc_vol_to_area | -0.56 | 0.00 |
| RHO GTPase Effectors | heightRange_mito.p | -0.56 | 0.00 |
| mRNA Splicing | sma_cell | -0.56 | 0.00 |
| Mitotic G1-G1/S phases | area_nucleus.p | -0.56 | 0.00 |
| Resolution of Sister Chromatid Cohesion | pixels_mito.p | -0.56 | 0.00 |
| G1/S Transition | area_cell | -0.56 | 0.00 |
| Extracellular matrix organization | nuc_vol_to_area | 0.56 | 0.00 |
| Cell Cycle, Mitotic | pixels_mito.p | -0.56 | 0.00 |
| Mitochondrial translation initiation | nuc_vol_to_area | -0.56 | 0.00 |
| Mitotic G2-G2/M phases | area_mito.p | -0.56 | 0.00 |

| | | | |
|---|---|---|---|
| Mitotic G2-G2/M phases | pixels_mito.p | -0.56 | 0.00 |
| GPCR ligand binding | pixels_cell | 0.56 | 0.00 |
| The citric acid (TCA) cycle and respirat... | count_cytoplasm.p | -0.56 | 0.00 |
| The citric acid (TCA) cycle and respirat... | count_mito.p | -0.56 | 0.00 |
| Mitotic Anaphase | csf_mito.p | -0.56 | 0.00 |
| Mitotic Anaphase | packing_mito.p | -0.56 | 0.00 |
| Class A/1 (Rhodopsin-like receptors) | sma_mito.p | 0.56 | 0.00 |
| Peptide ligand-binding receptors | sma_mito.p | 0.56 | 0.00 |
| Signaling by GPCR | area_nucleus.p | 0.56 | 0.00 |
| G1/S Transition | area_nucleus.p | -0.56 | 0.00 |
| Mitotic Metaphase and Anaphase | csf_mito.p | -0.56 | 0.00 |
| Mitotic Metaphase and Anaphase | packing_mito.p | -0.56 | 0.00 |
| Condensation of Prometaphase Chromosomes | pixel_per_mito_avg | -0.56 | 0.00 |
| G alpha (q) signalling events | pixels_cell | 0.56 | 0.00 |
| G2/M Transition | area_mito.p | -0.56 | 0.00 |
| Mitochondrial translation termination | area_cell | -0.56 | 0.00 |
| Signaling by GPCR | vol_mito.p | 0.56 | 0.00 |
| Cell Cycle Checkpoints | vol_mito.p | -0.56 | 0.00 |
| Class A/1 (Rhodopsin-like receptors) | sma_cell | 0.55 | 0.00 |

**Supplementary Table 1:** Correlation between imaging and sequencing derived feature pairs. Only the top 150 most significant correlations are displayed. P-values are Bonferroni-corrected.

| value | Description |
|---|---|
| vol_cytoplasm.p | volume of cytoplasm |
| vol_mito.p | volume of mitochondria |
| vol_nucleus.p | volume of nucleus |
| area_cytoplasm.p | area of cytoplasm |
| area_mito.p | area of mitochondria |
| area_nucleus.p | area of nucleus |
| pixels_cytoplasm.p | total pixel count of cytoplasm |
| pixels_mito.p | total pixel count of mitochondria |
| pixels_nucleus.p | total pixel count of nucleus |
| count_cytoplasm.p | number of cytoplasm DBSCAN clusters |
| count_mito.p | number of mitochondria DBSCAN clusters |
| z | slice depth |
| area_cell | area of cell |
| vol_cell | volume of cell |
| heightRange_cell | height range of cell |
| convexity_cell | relative area of cell lost to indentations |
| packing_cell | packing of cell |
| sphericity_cell | roundness of cell: spheres have a sphericity of 1 |
| sma_cell | sma cell |
| maxAreaSlice_cell | maximum area of available crossections of the cell |
| rugosity_cell | degree of wrinkling of the cell |
| pixels_cell | total pixel count of cell |
| heightRange_cytoplasm.p | height range of cytoplasm |
| convexity_cytoplasm.p | relative area of cytoplasm lost to indentations |
| packing_cytoplasm.p | packing of cytoplasm |

| | |
|---|---|
| sphericity_cytoplasm.p | roundness of cytoplasm: spheres have a sphericity of 1 |
| sma_cytoplasm.p | sma cytoplasm |
| csf_cytoplasm.p | csf cytoplasm |
| MinDist_cytoplasm.p_mito.p | Distance between cytoplasm and mitochondria (minimum) |
| MaxDist_cytoplasm.p_mito.p | Distance between cytoplasm and mitochondria (maximum) |
| MedianDist_cytoplasm.p_mito.p | Distance between cytoplasm and mitochondria (median) |
| MinDist_cytoplasm.p_nucleus.p | Distance between cytoplasm and nucleus (minimum) |
| MaxDist_cytoplasm.p_nucleus.p | Distance between cytoplasm and nucleus (maximum) |
| MedianDist_cytoplasm.p_nucleus.p | Distance between cytoplasm and nucleus (median) |
| heightRange_mito.p | height range of mitochondria |
| convexity_mito.p | relative area of mitochondria lost to indentations |
| packing_mito.p | packing of mitochondria |
| sphericity_mito.p | roundness of mitochondria: spheres have a sphericity of 1 |
| sma_mito.p | sma mitochondria |
| csf_mito.p | csf mitochondria |
| MinDist_mito.p_nucleus.p | Distance between mitochondria and nucleus (minimum) |
| MaxDist_mito.p_nucleus.p | Distance between mitochondria and nucleus (maximum) |
| MedianDist_mito.p_nucleus.p | Distance between mitochondria and nucleus (median) |
| heightRange_nucleus.p | height range of nucleus |
| convexity_nucleus.p | relative area of nucleus lost to indentations |
| packing_nucleus.p | packing of nucleus |
| sphericity_nucleus.p | roundness of nucleus: spheres have a sphericity of 1 |
| sma_nucleus.p | sma nucleus |
| maxAreaSlice_nucleus.p | maximum area of available crossections of the nucleus |
| rugosity_nucleus.p | degree of wrinkling of the nucleus |
| meanIntensity_cytoplasm.p | average intensity of cytoplasm |
| medianIntensity_cytoplasm.p | median intensity of cytoplasm |
| maxIntensity_cytoplasm.p | maximum intensity of cytoplasm |
| minIntensity_cytoplasm.p | minimum intensity of cytoplasm |
| meanIntensity_mito.p | average intensity of mitochondria |
| medianIntensity_mito.p | median intensity of mutochondria |
| maxIntensity_mito.p | maximum intensity of mitochondria |
| minIntensity_mito.p | minimum intensity of mitochondria |
| pixel_per_mito_avg | average pixel count per mitochondria |
| pixel_per_volume_mito | resolution of mitochondria |
| nuc_to_mito | ratio of nuclei count to mitochondrial count |
| pixels_per_volume_cytoplasm.p | resolution of cytoplasm |
| pixels_per_volume_mito.p | resolution of mutitochondria |
| pixels_per_volume_nucleus.p | resolution of nucleus |
| nuc_vol_to_area | ratio of nuclear volume to nuclear area |

**Supplementary Table 2:** Definition of imaging features

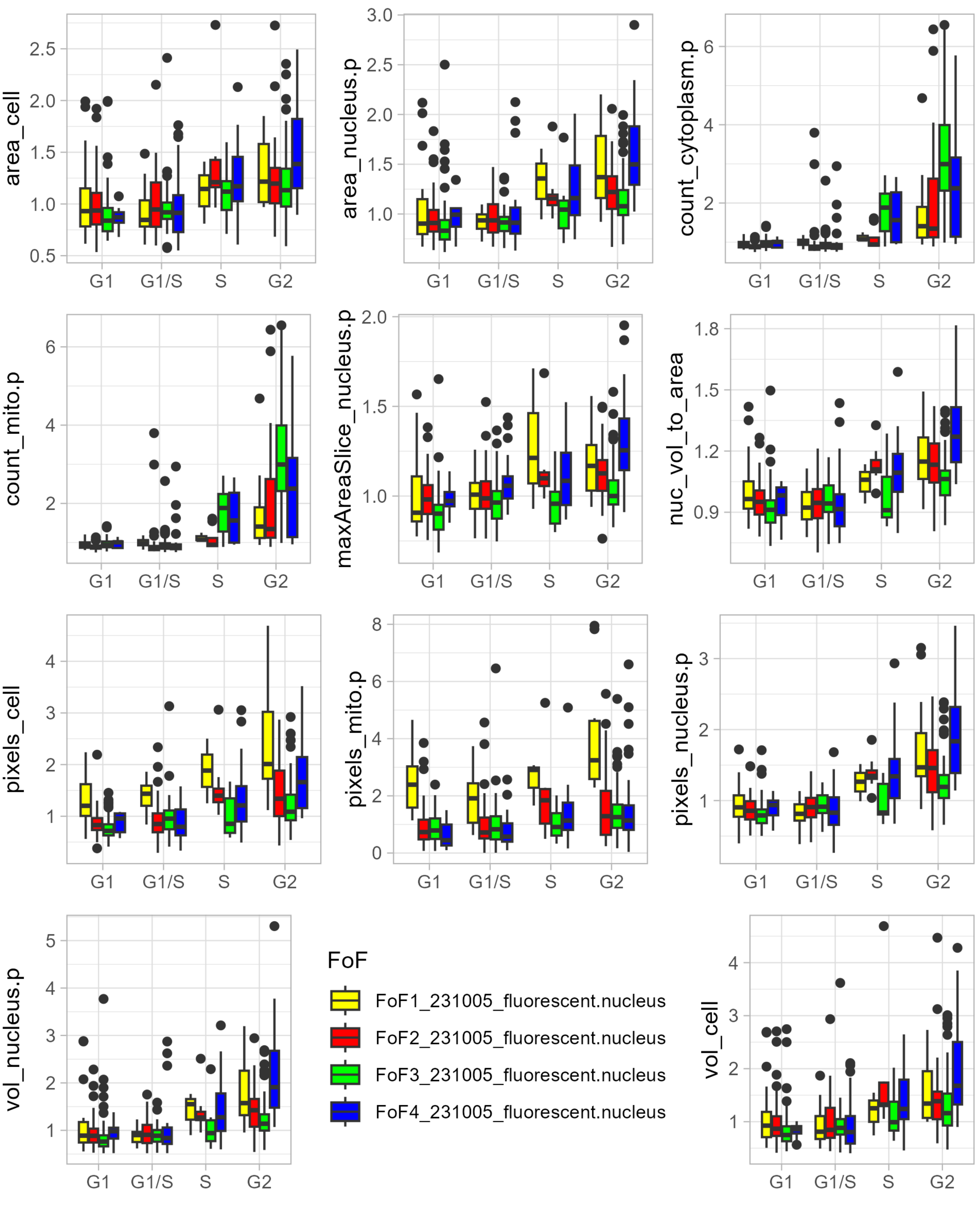


**Supplementary Figure 2:** Cell features derived from label free imaging correlate with FUCCI derived cell cycle progression. These features were prioritized based on their differential distribution across FUCCI-derived cell cycle classes (Anova test: p-value$<$ 0.01 across all four fields of view.)

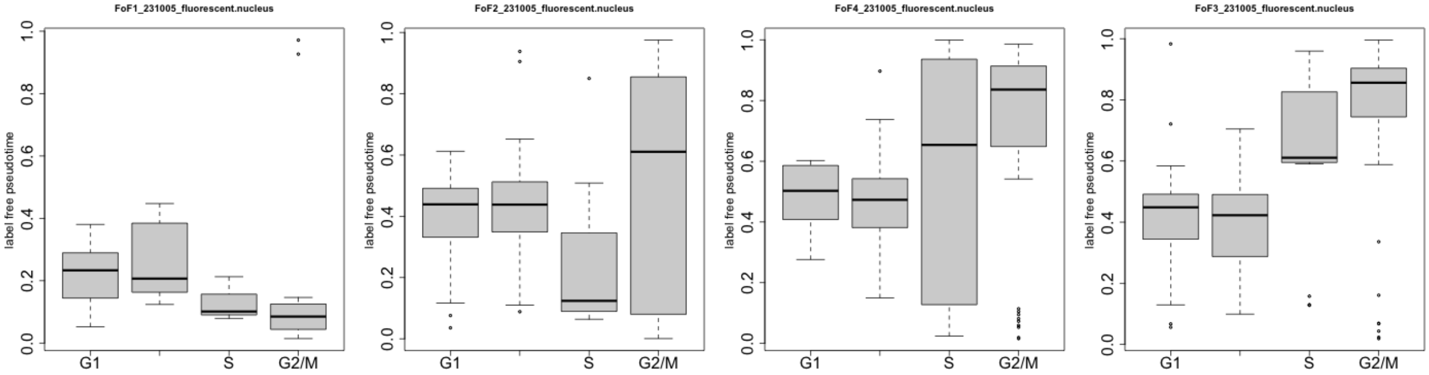


**Supplementary Figure 3:** Pseudtime derived from label-free imaging is differentially distributed across the four cell cycle phases. .

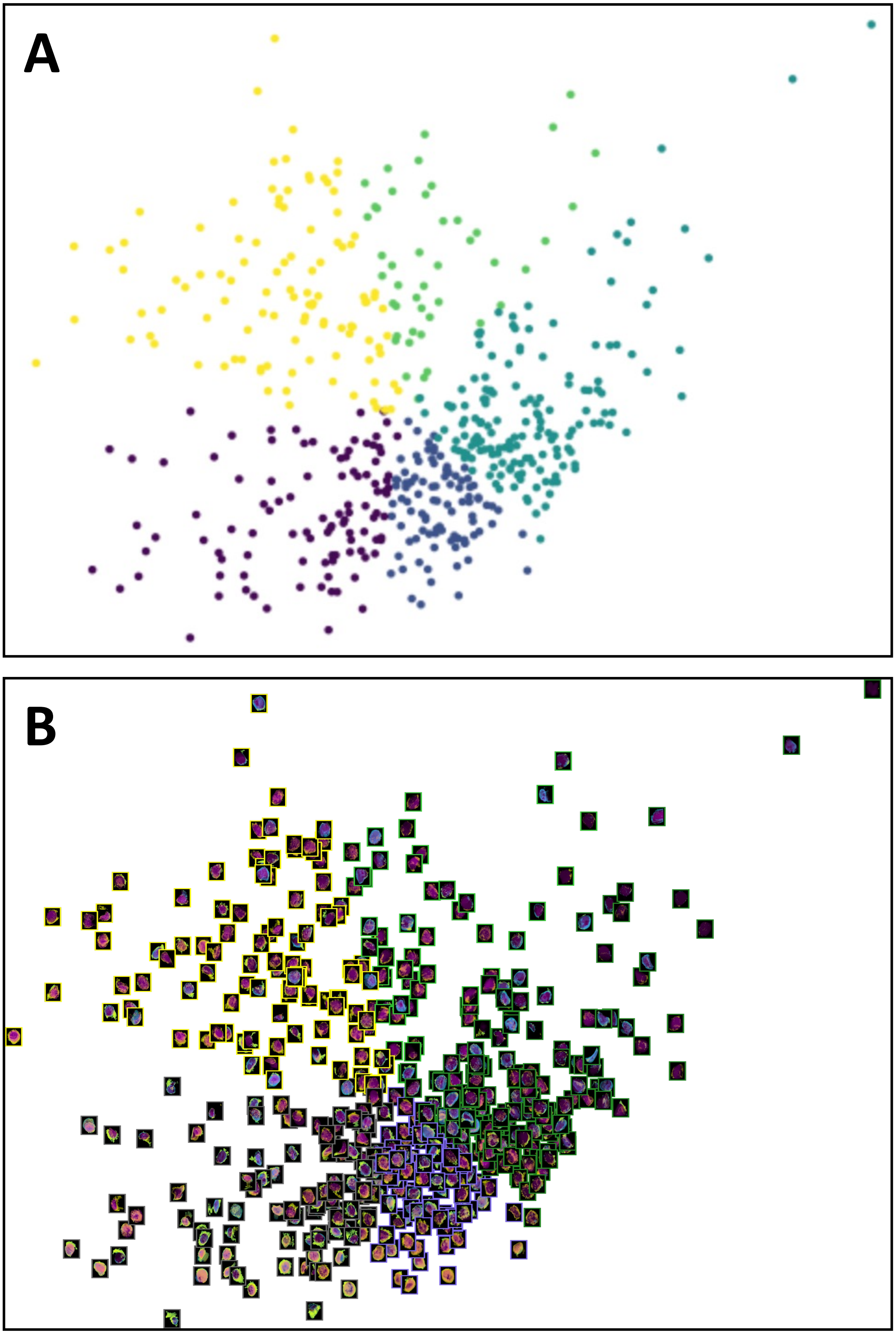


**Supplementary Figure 4:** Figure showing Image tSNE generated using Mistic for 506 single cell images [44]. Images are arranged based on their pseudo times and user-defined 2D tSNE coordinates. Each image has a border that matches the pseudotime it belongs to. Colorbar refers to that in Fig. 4C).

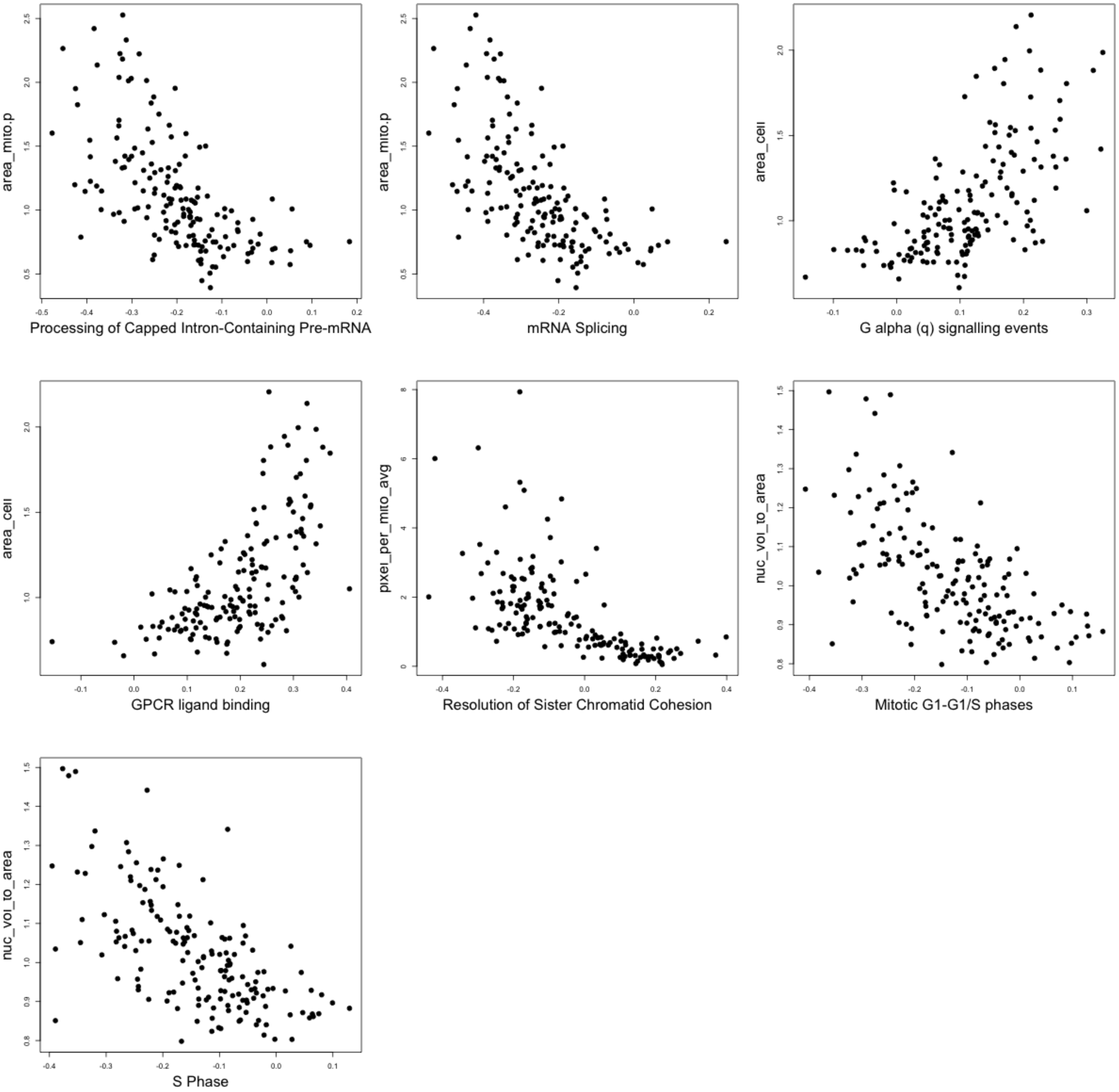


**Supplementary Figure 5:** Sequencing- and imaging derived features are correlated. .

### References


[1] U. Ben-David, B. Siranosian, G. Ha, H. Tang, Y. Oren, K. Hinohara, C. A. Strathdee, J. Dempster, N. J. Lyons, R. Burns, A. Nag, G. Kugener, B. Cimini, P. Tsvetkov, Y. E. Maruvka, R. O'Rourke, A. Garrity, A. A. Tubelli, P. Bandopadhayay, A. Tsherniak, F. Vazquez, B. Wong, C. Birger, M. Ghandi, A. R. Thorner, J. A. Bittker, M. Meyerson, G. Getz, R. Beroukhim, and T. R. Golub. Genetic and transcriptional evolution alters cancer cell line drug response. 560(7718):325–330. PMID: 30089904.

[31] J. Casale and J. S. Crane. Biochemistry, Glycosaminoglycans. In StatPearls. StatPearls Publishing.

[32] A. C. Lloyd. The Regulation of Cell Size. 154(6):1194–1205.

[33] P. Mishra and D. C. Chan. Mitochondrial dynamics and inheritance during cell division, development and disease. 15(10):634–646. PMID: 25237825.
